# Neuronal transcriptome analyses reveal novel neuropeptide modulators of excitation and inhibition imbalance in *C. elegans*

**DOI:** 10.1101/821769

**Authors:** Katherine A. McCulloch, Kingston Zhou, Yishi Jin

**Affiliations:** Section of Neurobiology, Division of Biological Sciences, University of California San Diego, La Jolla, California 92093

**Keywords:** *ins-29*, Insulin-like peptide, *acr-2*, *ets-5*, *flp-12*, RNA-seq, cholinergic excitation, locomotion

## Abstract

Neuropeptides are secreted molecules that have conserved roles modulating many processes, including mood, reproduction, and feeding. Dysregulation of neuropeptide signaling is also implicated in neurological disorders such as epilepsy. However, much is unknown about the mechanisms regulating specific neuropeptides to mediate behavior. Here, we report that the expression levels of dozens of neuropeptides are up-regulated in response to circuit activity imbalance in *C. elegans. acr-2* encodes a homolog of human nicotinic receptors, and functions in the cholinergic motoneurons. A hyperactive mutation, *acr-2(gf)*, causes an activity imbalance in the motor circuit. We performed cell-type specific transcriptomic analysis and identified genes differentially expressed in *acr-2(gf)*, compared to wild type. The most over-represented class of genes are neuropeptides, with insulin-like-peptides (ILPs) the most affected. Moreover, up-regulation of neuropeptides occurs in motoneurons, as well as sensory neurons. In particular, the induced expression of the ILP *ins-29* occurs in the BAG neurons, which are previously shown to function in gas-sensing. We also show that this up-regulation of *ins-29* in *acr-2(gf)* animals is activity-dependent. Our genetic and molecular analyses support cooperative effects for ILPs and other neuropeptides in promoting motor circuit activity in the *acr-2(gf)* background. Together, this data reveals that a major transcriptional response to motor circuit dysregulation is in up-regulation of multiple neuropeptides, and suggests that BAG sensory neurons can respond to intrinsic activity states to feedback on the motor circuit.

**AUTHOR SUMMARY:** Neuropeptides are secreted small molecules that regulate a variety of neuronal functions and are also implicated in many diseases. However, it remains poorly understood how expression of neuropeptides is regulated, particularly in disease states. Using a genetic animal model that mimics epilepsy, we identified dozens of neuropeptides that are up-regulated when neuronal activities are altered. Some of these neuropeptides share similarity to insulin-like properties (ILPs). Strikingly, one of these ILPs is expressed in sensory neurons that normally respond to acute carbon dioxide exposure. We show that the mis-regulation of this ILP expression is activity-dependent. Moreover, these neuropeptides act in concert to modulate animal behaviors. The findings in this study provide further evidence that neuropeptides are key mediators of aberrant cholinergic signaling, and suggest complex neural network effects from sensory neurons onto motor function.

## INTRODUCTION

Neural circuits are dynamic, changing their properties in response to experience. These changes are critical for maintaining circuit homeostasis and in processes like memory. Many factors such as c-Fos and BDNF, are activated early in response to increased neural activity and further regulate the expression of downstream genes [1]. These early-acting genes are also involved in many neurological diseases. For example, mutations in the activity-dependent transcriptional repressor gene Mecp2 are associated with Rett’s syndrome [2]. Mecp2 is necessary for the transcriptional up-regulation of BDNF, a key early-acting gene regulating synaptic plasticity [3].

Neuropeptides are small, secreted molecules that play neuro-modulatory roles in all animals. Neuropeptides have an extraordinarily diverse set of functions, including in feeding, mood, and reproduction, among others. Secreted neuropeptides bind to G-protein coupled receptors (GPCRs) on target cells to modulate neuronal activity. Neuropeptides can act on cells post-synaptic to where they are secreted from, but can also act over long distances. Neuropeptide expression can be changed by experience, for example, the expression of Neuropeptide Y(NPY) changes in response to a myriad of stressors. NPY can inhibit anxiety in multiple stress models [4], and may also play a role in neurological diseases, such as epilepsy [5]. Although multiple neuropeptides have been implicated in diseases, much remains unknown about how they are regulated [5,6].

The nematode *C. elegans* has long been an important experimental model for investigating the regulation of neuronal circuits. The well-defined connectomics of its nervous system, in combination with powerful genetics and molecular tools, enable *in vivo* dissection of neural circuit regulation with high resolution. *C. elegans* locomotion is controlled through the balanced activities of cholinergic excitatory neurons and GABAergic inhibitory neurons to promote contraction and relaxation of body-wall muscle, respectively. The locomotor circuit has been used to identify multiple conserved genes that regulate synaptic transmission. The gene *acr-2* encodes a neuronal acetylcholine receptor subunit that is expressed in cholinergic motor neurons. A Valine-to-Methionine transition mutation causes a gain-of-function in *acr-2* [*acr-2(gf*)] that results in a hyperactive channel [7]. The mutation affects a highly conserved residue within the pore-lining TM2 domain, and similar mutations in the human CHRNB2 cholinergic receptor subunit are associated with Autosomal Dominant Frontal Lobe Epilepsy (ADFLE) [8]. *acr-2(gf)* worms show defective movement as well as spontaneous whole-body shrinking, or convulsion.

Over 100 neuropeptide genes have been identified in *C. elegans*. These genes produce neuropeptides that fall into three classes: FMRFamide-like peptides (FLP), neuropeptide-like proteins (NLP) and insulin-like peptides (ILP) [9]. As with neuropeptides in humans, each neuropeptide gene produces a pro-neuropeptide, that is subjected to several enzymatic processing steps. The *flp* and *nlp* genes can generate several neuropeptides from a single locus through enzymatic cleavage by the pro-protein convertase *egl-3* and the endopeptidase *egl-21* [10,11]. Similar to human insulin and insulin-like growth factors, the *ins* loci produce a single peptide, that is activated from the pro-insulin peptide through the enzymatic activity of *egl-3* and a related pro-protein convertase *kpc-1* [12].

Previous work has shown that the neuropeptides *flp-18* and *flp-1* are important for regulating neurotransmission in *acr-2(gf)* mutants [13]. Additionally, *flp-18* expression was up-regulated in cholinergic motoneurons to inhibit convulsion of *acr-2(gf)* animals in a homeostatic manner. However, loss of function in the gene *unc-31*/CAPS, which is required for neuropeptide secretion, suppressed *acr-2(gf)* convulsion, suggesting that other neuropeptides function to promote circuit hyperactivity. Using a cell-type specific transcriptomic approach, we identified over 200 genes whose expression was significantly altered in *acr-2(gf)* neurons compared to wild type. Among them, genes involved in neuropeptide signaling were significantly over-represented in this gene list. One of these, *ins-29*, has not been previously characterized. Expression reporters for *ins-29* are weakly or not expressed in BAG gas-sensing neurons in wild type, and *ins-29* expression is clearly increased in *acr-2(gf)* animals. The increased *ins-29* expression in BAG neurons of *acr-2(gf)* adults is activity-dependent and requires the transcription factor *ets-5*, which has been shown previously to act embryonically to make BAG functionally competent [14]. Although BAG neurons interact with the motor circuit to respond to environmental cues, our data indicates that intrinsic activity states modulate expression of genes in the sensory BAG neuron, which then feed-back on the motor circuit. Finally, we present functional evidence supporting that the concerted action of several neuropeptides underlies motor circuit hyperactivity.

## RESULTS

### Expression profiling of adult cholinergic neurons

We performed cell-type specific RNA-seq analyses, using the *Pacr-2::gfp* reporter *juIs14* to isolate GFP expressing neuronal cells by FACS followed by RNA-seq (Materials and Methods, Table S1) [7,15,16,17]. Besides expression in cholinergic motoneurons (VA, VB, DA, and DB), GFP expressed from *juIs14* is also present in several unidentified neurons in the head and tail [7]. RNA-seq data from isolated wild-type neurons was analyzed with the Cufflinks program to identify expressed genes (Materials and Methods, Table S2). The gene list for the wild type cholinergic motor neurons was compared with a recent study that profiled major tissue-types in adult *C. elegans* [18]. We found that almost all of the genes identified in our dataset (∼95%) were also detected in a pan-neuronal analyses of expressed genes (Figure 1A), but shared very little overlap with tissue-specific expression profiles identifying enriched transcripts in hypodermis and muscle (Figure 1A). This comparison suggests our sample were relatively free of contamination from surrounding tissue. We also compared our data to previous analysis of a subset of cholinergic motoneurons (VA and DA) labeled with *Punc-4::gfp* [19]. Although the cell-types analyzed in these studies are not identical (A-type only), and they were performed at different life stages (larval vs. young adult in this study) using different techniques (microarray vs. RNA-seq in this study), over half of the genes (∼75%) from our dataset were also identified by microarray in larval type A motoneurons (Figure 1B). These include core cholinergic genes such as *unc-17*/VAChT (Avg. FPKM=484.1) and pan-neuronal genes such as *unc-13* (Avg. FPKM=90.9), required for synaptic vesicle priming [20,21]. Additionally, both of these datasets were relatively free of GABA specific transcripts. Thus, this comparison shows the genes identified in our sample are highly enriched for those that are validated to be expressed and function in cholinergic neurons.

**Figure 1.**
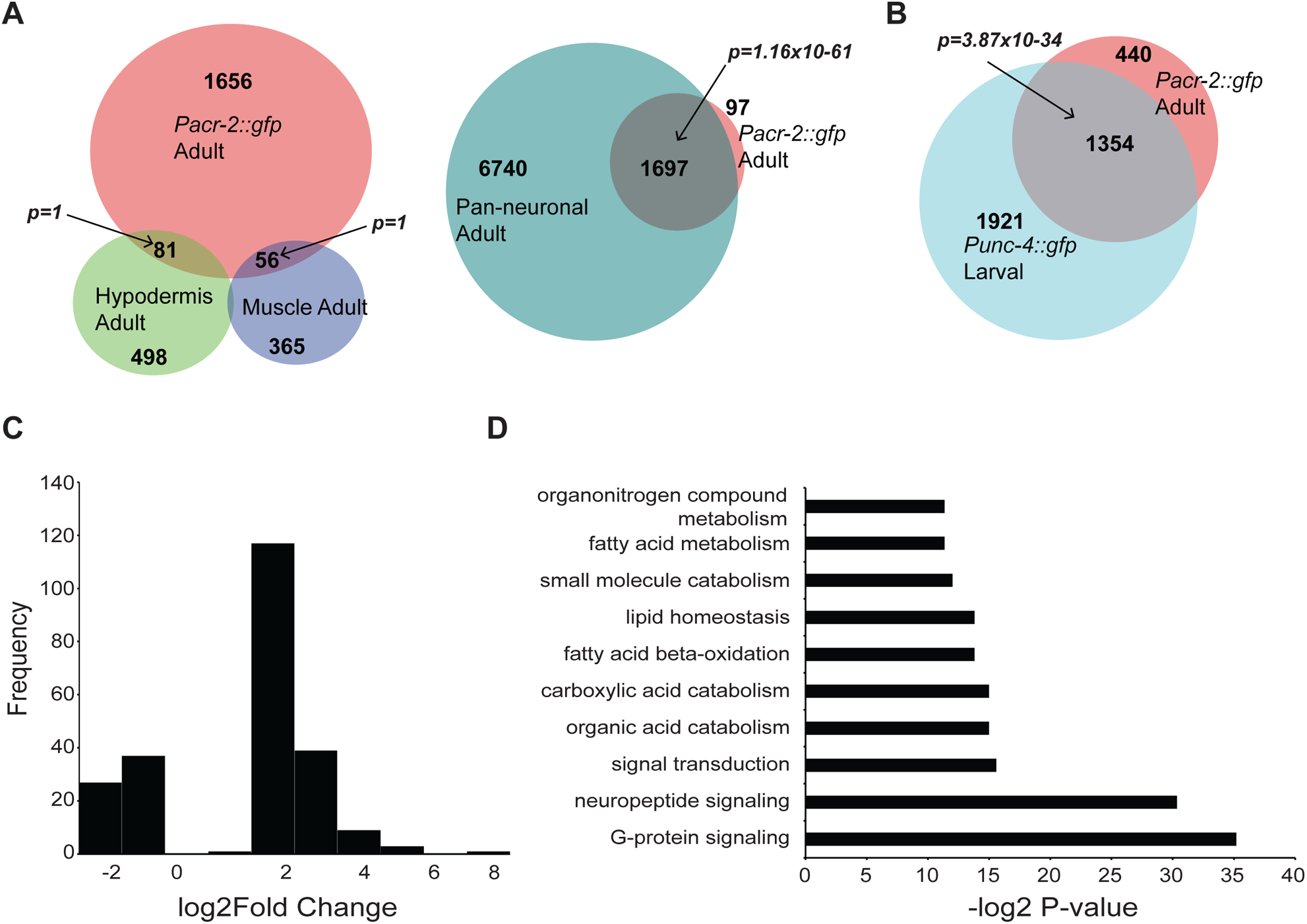
Differential expression analyses of *acr-2(gf)* neurons compared to wild type. A. Few genes identified as enriched in hypodermis or muscle are detected in isolated adult neurons labeled by *Pacr-2::gfp*. In contrast, almost all of the genes identified as expressed in cholinergic neurons labeled by *Pacr-2::gfp* are identified in pan-neuronal data-sets. Shown are Venn Diagrams with overlaps between the indicated data sets. The number in each region indicates the number of genes in that category. Hypodermis, muscle, and neuronal datasets are from Kaletsky *et. al*. (2018). Statistics for significance of overlap were performed using a hypergeometric distribution with the *phyper* function in R. B. Genes identified in previous expression profiling of larval cholinergic motor neurons, including core neuronal and cholinergic markers, are identified by RNA-seq of *Pacr-2::gfp* expressing neurons. The Venn Diagram displays that over half of genes identified by microarray as expressed in larval A-type motor neurons are also identified by RNA-seq in adult cholinergic neurons that express P*acr-2::gfp*. The number in each region indicates the number of genes in that category. Larval A-type motor neuron expression data is from Von Stetina *et. al*. (2007). Statistics for significance of overlap were performed using a hypergeometric distribution with the *phyper* function in R. C. Histogram of log_2_(Fold Change) for significantly different genes. Most genes that were different in *acr-2(gf)* compared to wild type were up-regulated at around 2-fold. D. GO-term analyses of genes significantly different in *acr-2(gf)* neurons compared to wild type (see Materials and Methods). Genes involved in neuropeptide and G-protein signaling (which included neuropeptide genes) were the most affected.

### Differential expression analyses between wild-type and *acr-2(gf)* neurons

We are interested in the genes differentially expressed in response to altered neuronal activity. The cholinergic neuron transcriptome in adult *acr-2(gf)* mutants was compared to those from wild type animals for changes in gene expression using DESeq2, and this analysis identified 234 genes as significantly mis-expressed in the *acr-2(gf)* mutant (Table S2) [22,23]. We found *flp-18* to be significantly up-regulated in *acr-2(gf)*, as predicated from our previous studies [13]. Analysis of the distribution of the fold change data using a histogram showed that a majority of the changes were up-regulation, at around 2-fold (Figure 1C). GO-term analyses indicated that genes involved in neuropeptide signaling were highly enriched in our gene list (Figure 1D) [24]. In total, the expression of 21 neuropeptide genes are identified as significantly up-regulated in *acr-2(gf)* neurons, representing approximately 18% of the estimated 113 neuropeptide genes identified in *C. elegans* (Figure 2A) [9]. Of these, several insulin-like peptides (ILP) were the most up-regulated (Figure 2A, Table S2). In addition, none of these ILP genes are detected as expressed in wild type cholinergic samples (Table S2). Together, these data indicate that the major response to altered motor circuit activity at the transcriptional level is to increase neuropeptide gene expression.

**Figure 2.**
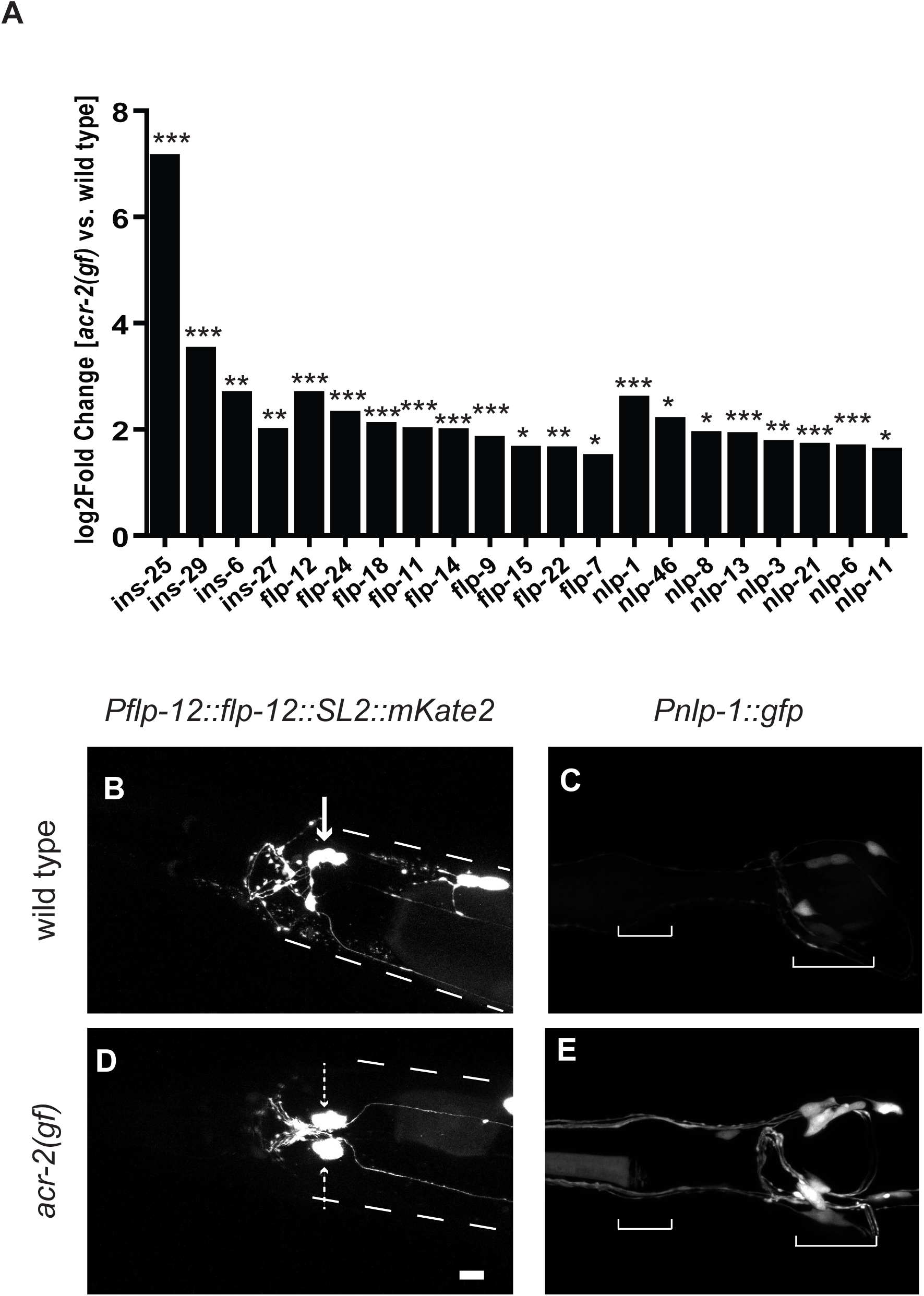
Neuropeptides are up-regulated in head neurons in *acr-2(gf)* animals. A. Log_2_(Fold Change) values for all up-regulated neuropeptides in *acr-2(gf)* animals is shown organized by class, as identified with DESeq2 analyses. Neuropeptides of each class were affected. (*P<0.05, **P<0.01, ***P<0.001) B-E. Differential expression of neuropeptide transcriptional reporters in the head in *acr-2(gf)* animals compared to wild type. (B,D)The arrow points to the likely SMB neuron expressing the *flp-12* reporter (*juEx7964*) in wild type based on shape and location. Dashed arrows point to the bilateral pair ectopically expressing the reporter in *acr-2(gf)*. Dashed lines indicated approximate outline of the animal. The posterior fluorescent signal in both strains in the co-injection marker labeling coelomocytes (C,E) *nlp-1::gfp (juEx7879)* expression is both increased in the same cells as wild type and also ectopically expressed. Brackets delineate the anterior and posterior pharyngeal bulbs, respectively.

To validate the RNA-seq analyses, we analyzed transcription reporters for the most up-regulated neuropeptide gene of each class. *flp-12* and *nlp-1* were the most up-regulated neuropeptides of their respective classes. These genes have been implicated in context-dependent behaviors. *flp-12* is involved in male locomotion, and *nlp-1* functions in food-evoked turning behaviors [25,26]. 2kb upstream of *flp-12* and *nlp-1* was used to generate GFP reporters, and both reporters showed expression in neurons in the head, similar to published studies of these neuropeptides (Figure 2B-E) [27,28]. In *acr-2(gf*), the *flp-12* reporter showed a different expression pattern than wild type, with ectopic expression in a pair of neurons close to the ganglion (Figure 2B,D). Similarly, the *nlp-1* reporter was up-regulated and expressed in additional cells in *acr-2(gf)* animals compared to wild type (Figure 2C,E). Therefore, this analysis provides an explanation for the increased RNA levels detected in RNA-seq of *acr-2(gf*), and also shows that increased locomotor excitation may cause mis-expression of these neuropeptide genes.

### *acr-2(gf)* increases expression of *ins-29* and *acr-2* in BAG sensory neurons

Among all up-regulated neuropeptides, *ins-25* and *ins-29* showed the most dramatic increase (Figure 2A). As the expression pattern and function of these ILPs is unknown, we investigated their expression patterns in further detail. *ins-25* and *ins-29* are within a less than 2.5kb region on Chromosome I, with *ins-29* being upstream of *ins-25*, in an operon, separated by 697bp intergenic sequence (Materials and Methods, Figure 3A). We used 2kb of sequence upstream of *ins-29* to drive GFP expression. In wild type animals, the *ins-29* reporter was sometimes weakly expressed in two neurons in the head, but often expressed in just one neuron or no GFP expression was detected (Figure 3B). However, in *acr-2(gf)* animals, consistent strong expression of GFP was observed in the same two head neurons (Figure 3C). These neurons are likely sensory, as they extend dendrites out to the nose of the animal.

**Figure 3.**
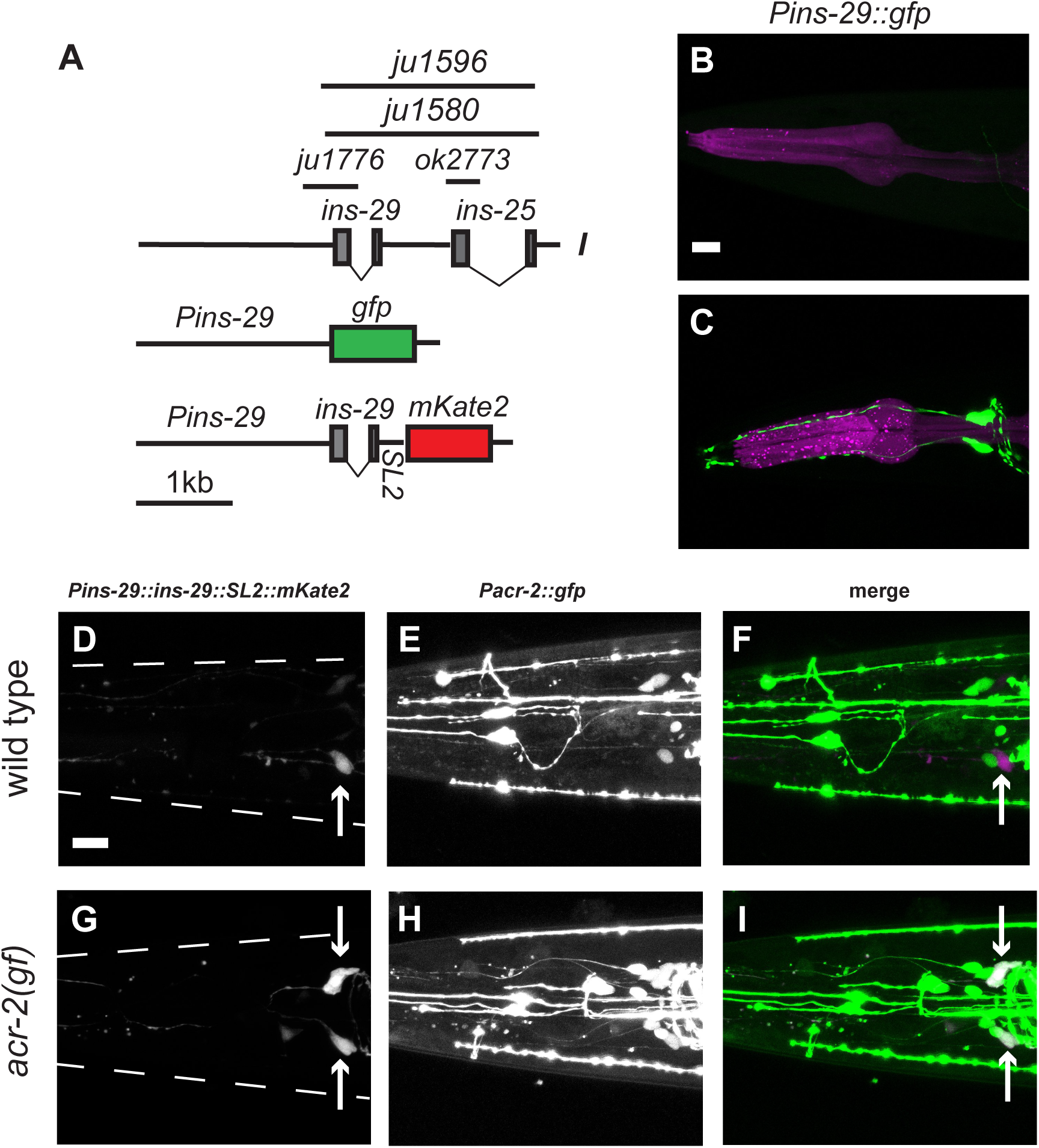
ILP gene and *acr-2* expression is up-regulated in two head neurons in *acr-2(gf)* A. Shown is the genomic region of *ins-29* and *ins-25* sequences. Insulin-like peptide genes all contain two exons, each encoding the A and B peptides, respectively. *ok2773* is a 354bp deletion in *ins-25. ju1776* is 568bp deletion in *ins-29. ju1580* is a 2.2kb deletion of both *ins-25* and *ins-29*, and a similar deletion *ju1596* is in *ins-27(ok2474)*, which is ∼6kb downstream of *ins-25*. Also shown are cartoons of the *ins-29* transcription reporters and expression constructs used in this study. SL2 sequence was inserted between *ins-29* and *mKate2* in the expression construct to monitor expression from the transgene without directly tagging the peptide. B-C. *Pins-29::gfp(juEx7742)* is strongly and consistently detected in two neurons in the head of *acr-2(gf)* animals, but often in just one or zero cells in wild type. Occasionally, *Pins-29::gfp* expression was observed in a third, more posterior neuron in both wild type and *acr-2(gf)*. D-I. A *Pins-29::ins-29::SL2::*mKate2 reporter*(juEx7966)* co-localizes with *Pacr-2::gfp(juIs14)* expression in *acr-2(gf*) animals, but not in wild type. scale bar=10µm. Cell bodies expressing mKate2 are labeled by an arrow. (D) In wild type animals, *Pins-29::ins-29::SL2::*mKate2(*juEx7966*) is weakly or not expressed in the head. Shown is an animal expressing the transgene in a single neuron. (E) *Pacr-2::gfp* is expressed in multiple neurons in the head of wild type animals. (F) Expression of *ins-29* and *acr-2* transcriptional reporters do not overlap in wild type animals, suggesting that *acr-2* is not normally expressed in the same neurons as *ins-29* in wild type. (G) Expression of *Pins-29::ins-29::SL2::mKate2* in *acr-2(gf)* is similar to that of *Pins-29::gfp* observed in (C). (H) Expression of *Pacr-2::gfp* reporter in *acr-2(gf)* animals. (I) Co-localization is observed between the *ins-29* reporter and *Pacr-2::gfp* in *acr-2(gf*) animals. In all images, animals are rolled to easily visualize neuron pairs. Beading observed in some images (i.e. the *Pacr-2::gfp*, (E),(H)), is due to rolling the animals in 10% agarose.

We next confirmed that the *ins-29* expression construct was expressed in cells labeled by *Pacr-2::gfp* (Figure 3B-E). We constructed an *ins-29* expression construct containing the endogenous *ins-29* trans-spliced to mKate2. The *Pins-29::ins-29::SL2::mKate2* expression pattern was similar to *Pins-29::gfp* (Figure 3D,E). We then generated strains co-expressing *Pins-29::ins-29::SL2::mKate2* and P*acr-2::gfp*. In wild-type, *Pacr-2::gfp* was expressed in several unidentified neurons in the head, in addition to its reported expression in cholinergic motoneurons [7]. No co-localization was observed between *Pacr-2::gfp* and the *Pins-29::ins-29::SL2::mKate2* in wild type animals that did express the *ins-29* reporter (Figure 3D-F). However, the *ins-29* expression construct showed co-localization with *Pacr-2::gfp* in *acr-2(gf)* animals (Figure 3G-I). This result suggests that, although total mRNA levels of *acr-2* in neurons are not significantly affected by the *acr-2(gf)* mutation, there is increased expression of *acr-2* in some head neurons.

The dendrite morphology and cell position of cells expressing *ins-29* suggested that they were likely to be the BAG neurons, which are known for their role in gas-sensing, particularly CO2 avoidance (Figure 3C) [29,30,31]. To confirm that the cells expressing *ins-29* were indeed BAG neurons, we generated animals co-expressing *Pins-29::ins-29::SL2::mKate* extrachromosomal arrays and an integrated GFP marker for BAG [*Pgcy-33::gfp*] and looked for co-localization (Figure 4A-F). We compared the expression of this construct to *Pgcy-33::gfp*, and, particularly in the *acr-2(gf)* background, where the *ins-29* reporter expression is consistently detected, the two expression patterns completely overlapped (Figure 4D-F). For those wild type animals that expressed the *ins-29* reporter, this expression also overlapped with the BAG marker (Figure 4A-C).

**Figure 4.**
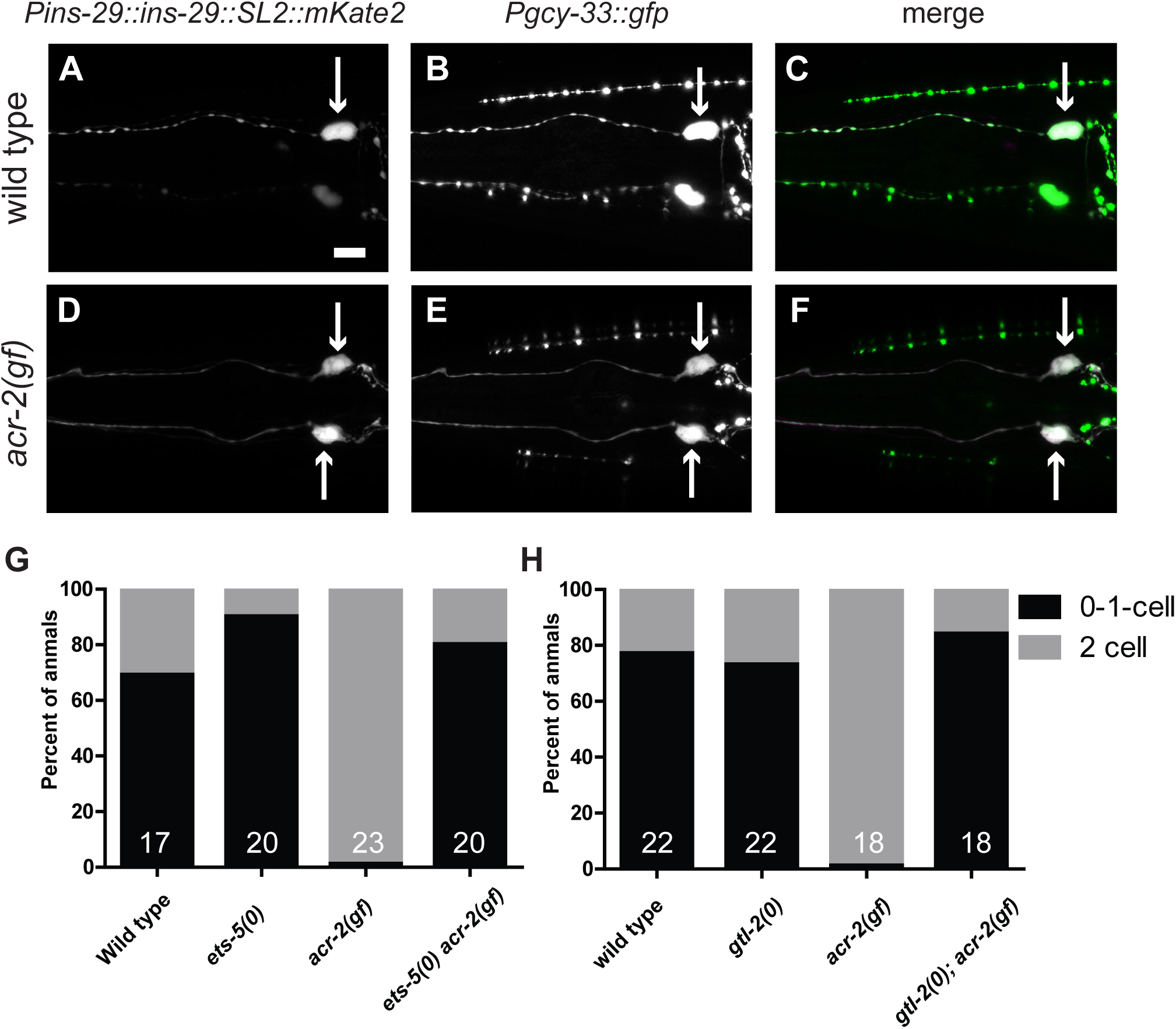
*ins-29* is expressed in BAG neurons and is regulated by the *ets-5* transcription factor and motor circuit activity. A-F. A *Pins-29* reporter is expressed in the BAG neurons. Cell bodies expressing indicated reporters are labeled by an arrow. (A) In wild type animals, expression of *Pins-29::ins-29::SL2::mKate2* in a single neuron in the head is shown. This reporter is often not expressed at all or in a single neuron in wild type, similar to the transcriptional GFP reporter (B) *Pgcy-33::gfp* labels BAG gas-sensing neurons in the head. (C) Overlap between the two reporters can be observed in one neuron. (D) Expression of *Pins-29::ins-29::SL2::mKate2* is observed in two head neurons, similar to the transcriptional GFP reporter. (E) Expression of *Pgcy-33::gfp* in *acr-2(gf)* is similar to wild type. (F) Complete overlap is observed between the two reporters. GFP-only signal is from the co-injection marker. Scale bar=10µm. In all images, animals are rolled to easily visualize neuron pairs. Beading observed in some images (i.e. the *Pgcy-33::gfp* and *Pmec-4::gfp* in B), is due to imaging plane of rolling the animals in 10% agarose. G. Almost all *acr-2(gf)* animals express *Pins-29::gfp(juEx7742)* in both BAG neurons, however mutation of *ets-5* causes animals to exhibit more similar *Pins-29::gfp* expression patterns as wild type. To assay *Pins-29::gfp* expression, animals were scored for detectable GFP in 0, 1, or 2 BAG neurons. N for each genotype is labeled in the bar. H. Mutation of the TRPM channel *gtl-2*, which almost completely suppresses *acr-2(gf)* convulsion and locomotion phenotypes, also restores *Pins-29::gfp(juEx7742)* expression to wild-type patterns. To assay *Pins-29::gfp* expression, animals were scored for detectable GFP in 0, 1, or 2 BAG neurons. N for each genotype is labeled in the bar.

The transcription factor *ets-5* is required for BAG development and function [14]. To determine if *ets-5* is involved in the expression of *ins-29* in *acr-2(gf)*, an *ets-5(0) acr-2(gf)* double mutant strain with the *Pins-29::gfp* was generated. Penetrance of expression was quantified by scoring for presence of GFP expression in zero, one, or both BAG neurons (Figure 3G). Very few wild-type animals express the transgene in both BAG neurons (31%), while most *acr-2(gf)* animals (90%) do. We found a marked decrease in the expression of the *ins-29* reporter in *acr-2(gf)* animals that lacked *ets-5* function, and these animals were more similar to wild type. Altogether, these data indicate that *ins-29* is up-regulated in sensory BAG neurons in an *ets-5*-dependent manner in response to *acr-2(gf)*.

### Reduction in circuit hyperactivity in *acr-2(gf)* animals restores *Pins-29::gfp* expression to wild type patterns

To further address whether the up-regulation of *Pins-29::gfp* in response to *acr-2(gf)* is dependent on neuronal activity, we next tested if reduction in motor circuit hyperactivity could restore expression of *ins-29* to wild type. The TRPM channel *gtl-2* is expressed in the hypodermis and regulates systemic ion homeostasis [32]. Mutation of this non-neuronal gene almost completely suppresses *acr-2(gf)* convulsion and locomotion phenotypes. The *ins-29* transcriptional *gfp* reporter was crossed into *gtl-2* null mutants in either a wild type or *acr-2(gf)* backgrounds. Animals were scored for GFP expression in zero, one, or both neurons (Figure 4H). *gtl-2(0)* animals alone resembled wild type. The expression pattern of *Pins-29::gfp* in *gtl-2(0); acr-2(gf)* animals also resembled wild type rather than *acr-2(gf)* alone. This result supports the idea that the changes in neuropeptide expression are likely due to systemic motor activity.

### Insulin-like peptides and *flp-12* coordinately promote motor circuit activity

Next, we addressed whether these neuropeptides expressed from head neurons affect motor circuit function. Genetic null [designated as *(0)*] mutations were used to determine if candidate up-regulated neuropeptides had functional roles in the *acr-2(gf)* motor circuit (see Materials and Methods). We focused on the most up-regulated neuropeptide genes of each class. Single loss of function mutations of *ins-25, ins-29, flp-12*, or *nlp-1* did not significantly affect convulsion rate (Figure 5A, S1). Additionally, a deletion mutation encompassing both *ins-29* and *ins-25* generated by CRISPR also did not significantly affect convulsion rate (Figure 3A, S1). However, a slight decrease in convulsion rate was observed in *flp-12(0) acr-2(gf)* animals (Figure 5A). Therefore, we made several combinations between *flp-12(0) acr-2(gf)* animals and deletion mutations in *ins-29 ins-25* and *flp-24*, the second most up-regulated *flp*-peptide gene. Analysis of convulsion rates in these strains showed that deletion of *flp-12* with either *flp-24* or *ins-29 ins-25* resulted in a significant decrease in convulsion rate compared to *acr-2(gf)* alone. This analysis suggests that these neuropeptides act to promote circuit hyperactivity.

**Figure 5.**
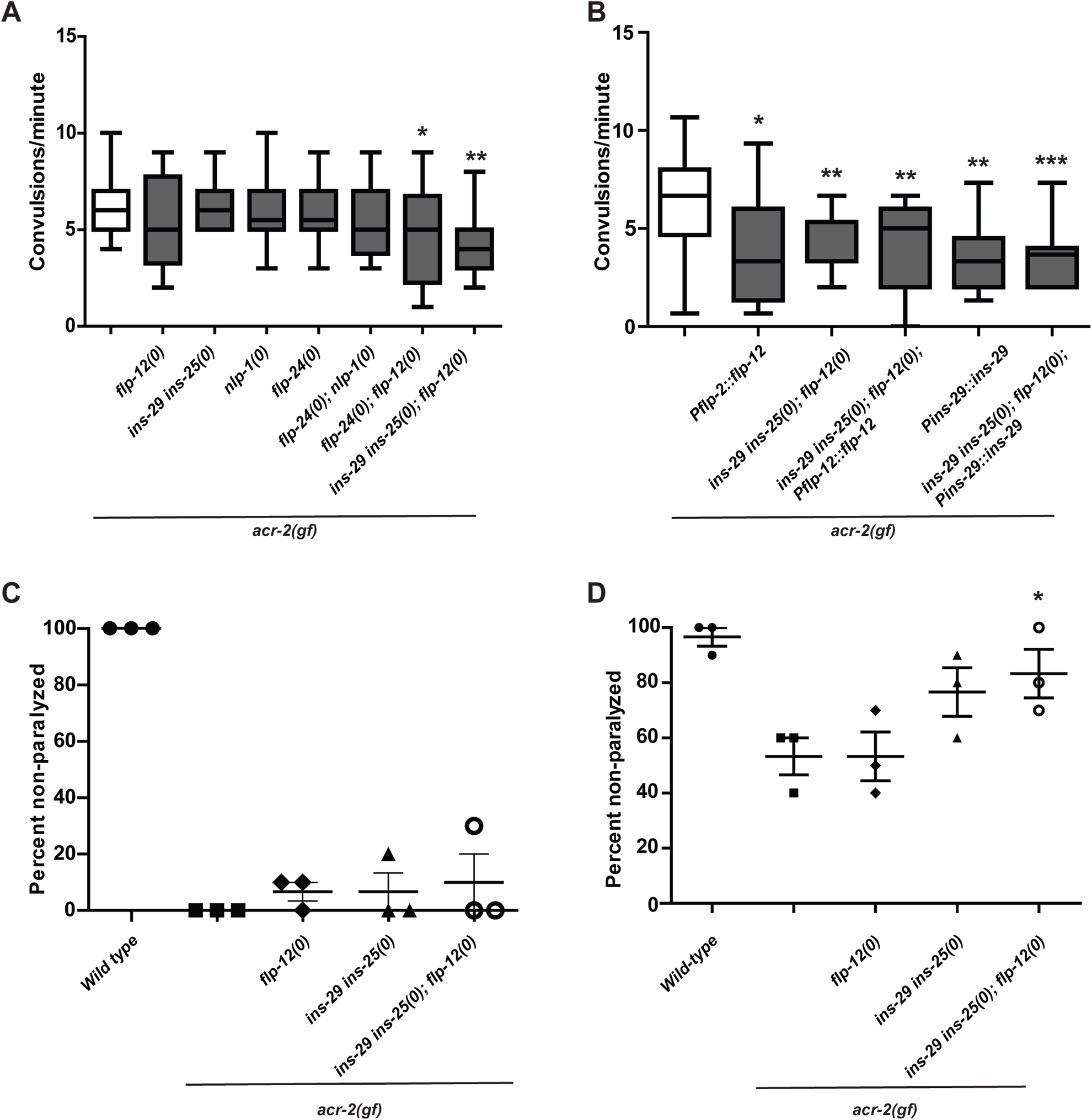
Insulin-like peptides and *flp-12* coordinately promote motor circuit activity. A. Convulsion rates for indicated genotypes are shown as convulsions/minute. Deletion of *flp-12* in combination with either *flp-24(0)* or deletion of the *ins-29 ins-25* region causes a statistically significant reduction in convulsion rate. (*P<0.05, **P<0.01, n.s.=non-significant. One-way ANOVA followed by Dunnett’s test. N≥19) B. Over-expression of either *flp-12* or *ins-29* suppresses convulsion. (*P<0.05, **P<0.01,**P<0.001, n.s.=non-significant. One-way ANOVA followed by Dunnett’s test. N≥19) C. Aldicarb sensitivity shown as percentage of animals not paralyzed after 1hour on the drug (Data from Figure 4A). No significant difference was observed. (Two-way ANOVA followed by Dunnett’s test, compared to *acr-2(gf)* alone.). Data is also shown in Table S3. D. Levamisole Sensitivity of neuropeptide mutants in the *acr-2(gf)* background at 15min (Data from Figure 4C). Mutation of insulin-like peptides with *flp-12* significantly reduces levamisole sensitivity compared to *acr-2(gf)* at this timepoint. (*P<0.05, Two-way ANOVA followed by Dunnett’s test compared to *acr-2(gf)* single mutant.). Data is also shown in Table S5.

We also sought to restore convulsion frequency in *ins-29(0) ins-25(0); flp-12 (0) acr-2(gf)* compound mutants by over-expressing wild type *ins-29* or *flp-12* in *ins-29 ins-25(0); flp-12(0) acr-2(gf)* mutant animals. Interestingly, over-expression of either *ins-29* or *flp-12* alone also partially suppressed convulsion frequency of *acr-2(gf)*, and the phenotype was similar in the neuropeptide compound mutant background (Figure 5B). Therefore, over-expression of either *ins-29* or *flp-12* acts in a dominant-negative manner to suppress *acr-2(gf)*. One possibility is that over-expression of these peptides blocks wild type function of their downstream receptor(s).

We further asked whether these neuropeptide genes affected synaptic transmission in the locomotor circuit using pharmacological assays. Aldicarb is an acetylcholinesterase inhibitor which leads to buildup of acetylcholine at the synaptic cleft and eventual paralysis in wild type animals [33]. Mutants that increase or decrease cholinergic activity display increased or decreased sensitivity to aldicarb. *acr-2(gf)* animals are more sensitive to aldicarb, consistent with their hyperactivity as described by previous electrophysiology and pharmacology analysis [7]. *ins-29(0) ins-25(0); flp-12(0) acr-2(gf)* animals were similar to *acr-2(gf)* alone on aldicarb, indicating that overall neurotransmission was not strongly affected by these mutations (Figure 5C, Table S3). *ins-29(0) ins-25(0); flp-12(0)* mutants were also not significantly different from wild type animals on aldicarb (Table S4). Levamisole is an agonist for the post-synaptic cholinergic receptors on muscle [34,35]. Resistance or sensitivity to levamisole can represent changes in muscle responsiveness. In these experiments, we found that *ins-29 ins-25(0); flp-12(0) acr-2(gf)* animals were less sensitive to levamisole than *acr-2(gf)* alone (Figure 5D, Table S5). However, no effect was observed in these compound neuropeptide mutants outside of the *acr-2(gf)* background (Table S6). This result indicates that these neuropeptides may function post-synaptically to affect motor circuit function, and this function is only observed in the *acr-2(gf)* background, consistent with these peptides being differentially expressed in response to *acr-2(gf)* hyperactivity. Together, these data show that insulin-like peptides and *flp-12* act together to promote *acr-2(gf)* circuit hyper-activity, at least partially by modulating post-synaptic function.

## DISCUSSION

Using neuronal-type specific RNA-seq, we identified over 200 genes whose expression levels are altered in response to cholinergic hyperactivity in the motor circuit of *C. elegans*. Genes encoding neuropeptides were over-represented in this list, and we validated that fluorescent reporters for *ins-29, flp-12*, and *nlp-1* were over-and/or ectopically-expressed in *acr-2(gf)* animals compared to wild type. These data support the conclusion that the major transcriptional response to cholinergic hyperactivity in *C. elegans* is by altering neuropeptide gene expression. Bioinformatic analyses of upstream sequences did not identify common motifs in the promoter regions of theses neuropeptides, suggesting that multiple transcriptional pathways may be involved in these changes. Indeed, the fluorescent reporters analyzed here showed diverse expression patterns as well as varied changes in expression, from over-expression in the same cells as wild type, to ectopic expression patterns.

Insulin-like peptides were the most up-regulated genes identified by RNA-seq in *acr-2(gf)*. The expression and function of one of these, *ins-29*, has not been previously characterized. Examination of *ins-29* transcriptional reporters showed expression in the gas-sensing BAG neurons, with increased intensity and more penetrant expression in both neurons detected in *acr-2(gf)* compared to wild type. Furthermore, co-labeling also showed up-regulation of *acr-2* itself in these neurons. BAG neurons are necessary for *C. elegans* to respond to changes in environmental CO2 [29]. The observation that *ins-29* is expressed in the BAG sensory neurons, rather than motoneurons or pre-motor interneurons, was surprising. For example, whereas *flp-18* is expressed from the cholinergic motoneurons to affect *acr-2(gf)* activity and motor function, *ins-29* expression is increased in sensory neurons in the head. Together, these results indicate that genes involved in cholinergic neurotransmission including the *acr-2* cholinergic receptor subunit gene, as well as the insulin-like peptide gene *ins-29* are up-regulated in BAG neurons in the *acr-2(gf)* background. Neuropeptide signaling from BAG has been shown previously to interact with another cholinergic circuit, the egg-laying circuit [36]. *flp-17* and *flp-10* secreted from BAG act in parallel with cholinergic signaling to inhibit egg-laying. It is proposed that signaling from BAG integrates favorable environmental signals with the egg-laying circuit.

The BAG-specific transcription factor *ets-5* also regulates *Pins-29::gfp* expression. Analysis of the *ins-29* promoter did not reveal a clear *ets-5* biding site, and it is possible this effect in indirect. *ets-5*-dependent pathways, therefore, are necessary for the increased expression of P*ins-29*::*gfp* in response to *acr-2(gf)*. This result also indicates a function for *ets-5* beyond development and maintenance of BAG neuron identity, but in modifying transcription in the BAG neurons under different physiological conditions in mature animals. Finally, loss of function or over-expression of ILP genes, along with *flp-12*, can suppress *acr-2(gf)* convulsion. Therefore, levels of multiple neuropeptides, such as the ILP INS-29 from BAG, can affect activity balances in the *C. elegans* motor circuit.

In humans, seizure is a common co-morbidity of diabetes. *In vitro* analysis has also found, for example, that IGF signaling can be neuroprotective in response to injury, but can also promote epileptogenesis[37]. Insulin peptide signaling may also play a role in Alzheimer’s Disease (AD) [6]. Although the role for insulin signaling in the brain is still unclear, levels of insulin and insulin receptors are markedly decreased in the brains of AD patients. Our data show that in *C. elegans*, ILP signaling is altered by aberrant cholinergic activity to modulate circuit function.

## MATERIALS & METHODS

### *C. elegans* genetics

*C. elegans* strains were maintained at 20-22°C. For RNA-seq experiments, CZ631(*juIs14[Pacr-2::gfp*]) and CZ5808 (*juIs14[Pacr-2::gfp]; acr-2(n2420)*)were used. For a list of all strains, see Table S1. All genetic null alleles are designated by *(0)* in the text.

CRISPR mutagenesis was performed as described previously, via co-CRISPR with *dpy-10* marker [38]. Two sgRNAs were designed to bind outside the *ins-29* and *ins-25* region [*ins-29* TTGGCGCCCAGCGCCGTTGT GGG, *ins-25* CAGATCTTCGATTGGGACGG CGG]. To make the *ins-29* single deletion, the 5’ *ins-29* sgRNA was injected. This generated a 568bp deletion spanning the entire first exon of *ins-29* (*ju1776*). Both sgRNAs were injected into wild-type or *ins-27(ok2474)* animals to delete both *ins-29* and *ins-25* and make an *ins-29 ins-25(0)* double mutant (*ju1580*) or *ins-29 ins-25(0) ins-27(0)* triple mutant chromosome (*ju1596 ok2474*). A ∼2.2kb mutation was isolated in each background and both alleles were crossed into *acr-2(gf)*. Most crosses (except those described below) were done using standard methods. See Table S7 for genotyping primers.

Construction of double mutant strains with *flp-12(ok2309)* or *ets-5(tm866)* with *acr-2(gf)* was made using MT6448 *lon-2(e678) acr-2(gf)* strain. *flp-12* (X:-7.20) and *ets-5* (X:-6.20) are to the left of *acr-2* (X:-2.56) on the X chromosome. We generated heterozygotes of the genotype: *lon-2(e678) acr-2(gf)* X/[*flp-12(ok2309)* or *ets-5(tm866)*] X. Non-long, convulsing progeny were isolated from the next generation and genotyped for the gene of interest.

### Sample preparation for RNA-seq

Samples were prepared for dissociation and FACS essentially as described [15,39]. Animals were synchronized by hypochlorite treatment. Eggs were isolated from wild type N2, CZ631, and CZ5808 gravid adults and allowed to hatch overnight and arrest at the L1-stage. Synchronized L1s were plated onto 15cm NGM plates seeded with OP50 bacteria. Approximately 8-10,000 L1s were plated to 15-20 plates for each strain. As N2 cells are only needed to control for the FACS sort, only 5 plates were prepared. These plates were incubated at 20°C for 72hr to reach adulthood prior to collection.

The entire process of cell preparation and sort was completed in a single day. Animals were washed off plates and spun down and the washed in M9 media to remove bacteria. Typically, pellets for each strain would be ∼500µl in volume, and these would be split into two tubes. 750µl of lysis buffer (200µM DTT, 0.25% SDS, 20mM HEPES pH8.0, 3% sucrose) was added to each tube and samples were incubated for approximately 6-7 minutes. Worms were then rapidly washed in M9 five times. Next, 500µl of 20mg/ml freshly made pronase solution was added. Samples were incubated in pronase approximately 20 minutes. Every 2-3 minutes, each sample was disrupted by pipetting with a P200, and samples were monitored for dissociation with a dissection microscope. When large worm chunks were no longer visible under the dissection microscope, the reaction was stopped by adding 250µl ice cold PBS-FBS solution (1X PBS solution with 2% Fetal Bovine Serum). Samples were then centrifuged in the cold at top speed in a microcentrifuge for 10 minutes, and pelleted cells were resuspended in 500µl FBS. Cells were then syringe-filtered (5µm pore). Samples were spun again in the cold and resuspended in ½ the starting volume of FBS. 80,000-100,000 GFP+ cells were collected for each sample at the UCSD Flow Cytometric Core in Moore’s Cancer Center. Cells were sorted directly into Trizol LS and stored at −80°C until preparation. RNA was isolated using Qiagen RNAeasy kit.

### RNA-seq analyses

RNA library preparation and sequencing were performed at the Institute for Genomic Medicine at UCSD. RNA libraries were prepared with Illumina TruSeq kit. Libraries were sequenced on an Illumina HiSeq4000. Data analyses were performed using the Galaxy platform [23]. We obtained 50-70 million single-end reads/sample. Reads for each sample were mapped and aligned using TopHat [40]. Identification of expressed genes in wild type was determined using Cufflinks. “Expressed” genes were selected by filtering for genes with an FPKM >10 in both replicates. This analysis produced 1,812 transcripts (Table S2). Using the BioVENN site, our list of “expressed genes” from wild type neurons was compared with those identified as enriched in adult epidermis and muscle and expressed in neurons by RNA-seq, as well as larval A-type motoneurons by microarray [18,19,41]. For differential expression analyses between wild type and *acr-2(gf*) neurons, BAM files were loaded into the HTSEQ program and then analyzed by DESeq2 for differential expression analyses (Table S2) [22,42]. GO-term enrichment analysis was performed using the GOrilla algorithm for enrichment of Biological Process terms[24]. Data analyses were performed in Microsoft Excel, R, and Graphpad Prism. Sequencing datasets have been deposited in the Gene Expression Omnibus (Accession GSE139212).

### Convulsion and pharmacological analyses

All behavior observations reported here were made on mutations that were outcrossed with N2 for at least 4 times. Convulsions were defined as simultaneous contraction of the body wall muscles producing a rapid, concerted shortening in body length. The convulsion frequency for day-1 adult animals was calculated during a 90-second period of observation. For levamisole sensitivity, ten day-1 adult animals were transferred to fresh plates containing 1 mM levamisole. After 1 hr, animals were assessed for paralysis; if plates contained non-paralyzed animals, then the strain was considered resistant to levamisole. Sensitivity to 500 μM or 1mM aldicarb was assessed by transferring ten day-1 adults to fresh aldicarb plates and by monitoring worms for paralysis every 30 minutes by gently touching the animal with a platinum wire. Aldicarb sensitivity was quantified for at least three independent experiments.

### Imaging and microscopy

Images of fluorescent reporter lines were taken on a Zeiss LSM 700 or 800 confocal microscope using the 63x objective. Animals were mounted in thick agarose (10%) and rolled with the ventral side up for consistent imaging. Hyperstacks were processed in ImageJ. (For images in Fig. 4, *Pgcy-33::gfp* strains, gains were set at 500V for mKate2 and 500V for GFP. For images in Fig. 3, of *juIs14* strains, gains were set at 550V for mKate2 and 600V for GFP.). All images are taken using the 63X objective. Scoring of *Pins-29::gfp* expression in Figure 4 was performed using a Zeiss Axioplan 2 at the 63X objective. A neuron was scored as “expressed” if GFP signal was clearly visible in the cell body and dendrites through the eyepiece.

### Molecular biology and *C. elegans* transformation

Transcriptional gfp reporters were made using Gibson assembly into pPD95.75. Vector was cut using restriction enzymes and PCR-amplified promoters were inserted. Approximately 2kb upstream of each neuropeptide gene was used as putative promoter sequence. Primers used for *Pins-29* were (gene-specific sequence in uppercase): Forward 5’tgcatgcctgcaggtcgactCTTTAAAATGGTTAATTTTGTAGTTAG3’/Reverse 5’tggccaatcccggggatcctTTTTTTATTTCACAATATAATATACTTTATAC3’. Primers used for *Pnlp-1* were: Forward 5’tgcatgcctgcaggtcgactTTGTTTTATCCAACATTATTCAC3’/Reverse 5’tggccaatcccggggatcctCGTTGCCTCAAGTTGATG3’. To generate mKate2 reporters for neuropeptide expression constructs, GFP sequence in pPD95.75 was replaced with SL2-mKate2 sequence. Genomic sequences for *flp-12* or *ins-29* was then amplified from genomic DNA and inserted into the modified pPD95.75 vector using Gibson Assembly. Primers for *ins-29*: Forward 5’aagcttgcatgcctgcaggtCTTTAAAATGGTTAATTTTGTAGTTAG3’/Reverse 5’tgaaagtaggatgagacagcTCAAGCAAGATTTGAAGG3’. Primers for *flp-12* were: Forward 5’tgcatgcctgcaggtACAACAAAAGTATTTTTGACG3’/ Reverse 5’agtaggatgagacagcCTACTTTCGTCCAAATCG3’. These sequences include the entire coding sequence plus 2kb upstream promoter.

cDNAs were generated using the SuperScript RT kit from Invitrogen. 2µg input RNA (from synchronized young adult *acr-2(gf)* populations) was used for each RT reaction. RNA was extracted and isolated using Trizol reagent and chloroform extraction. 1µl of RT (∼100ng) was used in each PCR reaction. Nested reactions were used for amplifying both *ins-29* and *ins-25* cDNAs. For *ins-29*, the first reaction used either SL1 or SL2 forward primer with a gene specific reverse primer: 5’ gcaagatttgaaggacagcac 3’. In the second reaction 2µl of the first PCR was used with Primers: Forward 5’TTCTGTAAATTTGTATTCCTGATC, Reverse 5’ GATTTGAAGGACAGCACAAT 3’. For *ins-25*, the first reaction used either SL1 or SL2 forward primers with a gene specific reverse primer :5’ caaatttgggcaacacatattc 3’. In the second reaction, 2µl of the first PCR reaction was used with Primers: Forward 5’ ATGTTGTTCAAAATCATCATT 3’, Reverse 5’ GGGCAACACATATTCTTCAG 3’. Product using the SL2 primer was only detected for ins-25 transcript and verified by Sanger sequencing.

*C. elegans* transgenic multi-copy arrays were generated using standard protocols [See Table S1 for list of transgenic strains made in this study] [43]. DNA was typically injected at 25ng/µl concentration. *flp-12* over-expression constructs were injected at 5ng/µl, as injection at 25ng/µl failed to yield transgenics.

## Acknowledgements

We thank our lab members for helpful discussions, and Matt Andrusiak and Ngang Heok Tang for comments on the manuscript. We also thank Rachel Kaletsky and Coleen Murphy for sharing advice on neuron isolation, and Martin Hudson for the XA2260 strain. Some strains were provided by the Caenorhabditis Genetics Center, which is funded by NIH Office of Research Infrastructure Programs (P40 OD010440) and the National Bioresource Project of Japan. K.A.M. was a trainee on NIH institutional training grants (T32 NS007220 and T32 AG000216). This work was supported by a NIH grant to Y. J. (R37 NS035546).

## SUPPORTING INFORMATION

**Figure S1. Analysis of insulin-like peptides in the motor circuit**

Convulsion rate of *acr-2(gf)* combined with different combinations of deletions in genes for insulin-like peptides up-regulated in *acr-2(gf)* neurons. None of these combinations had a statistically significant effect of convulsion rate (One-way ANOVA followed by Dunnett’s test).

**Table S1. Strains used in this study**

**Table S2. Transcriptome analyses of wild type and *acr-2(gf)* neurons**

**Table S3. Aldicarb timecourse for neuropeptide mutants with *acr-2(gf)***

**Table S4. Aldicarb timecourse for neuropeptide mutants**

**Table S5. Levamisole timecourse for neuropeptide mutants with *acr-2(gf)***

**Table S6. Levamisole timecourse for neuropeptide mutants**

**Table S7. Genotyping primers**

